# Exploring Conformational Transitions of RNA Dimers via Machine Learning Potentials

**DOI:** 10.64898/2026.02.25.707885

**Authors:** Leonardo Medrano Sandonas, Macarena Tolmos Nehme, Luis Fernando Cofas-Vargas, Gustavo E. Olivos-Ramirez, Gianaurelio Cuniberti, Simón Poblete, Adolfo B. Poma

**Affiliations:** Institute for Materials Science and Max Bergmann Center of Biomaterials, TUD Dresden University of Technology, 01062, Dresden, Germany; Universidad Nacional de Ingeniería, Av. Túpac Amaru 210, Rímac, Lima 15333, Peru; Departamento de Química, Universidad Autónoma Metropolitana-Iztapalapa, Mexico City C.P. 09310, Mexico; Department of Biosystems and Soft Matter, Institute of Fundamental Technological Research, Polish Academy of Sciences, ul. Pawińskiego 5B, 02-106, Warsaw, Poland; Cluster of Excellence CARE, TU Dresden and RWTH Aachen, Germany; Cluster of Excellence CeTI, TU Dresden, Germany; Facultad de Ingeniería, Universidad San Sebastián, Santiago, Chile; Centro BASAL Ciencia & Vida, Universidad San Sebastián, Santiago, Chile

## Abstract

RNA is a flexible biopolymer that adopts diverse conformations while forming structural motifs essential for its function. Classical RNA force fields often show limited transferability and inefficient sampling of transitions between stable states, particularly in moderately large RNA. To address these limitations, quantum-informed machine learning (ML) potentials have recently emerged as a promising alternative, offering improved accuracy and transferability relative to classical force fields. Here, we assess ML potentials for exploring RNA conformations using the adenine–adenine dinucleoside monophosphate (ApA) dimer, a fundamental RNA building block. We generated an extensive quantum-mechanical (QM) dataset for ApA conformations obtained from temperature replica exchange molecular dynamics (TREMD) simulations. Despite its small size, the ApA dimer exhibits six conformations in which quantum effects and solvent-mediated interactions play crucial roles. Using this dataset, we parameterized ML potentials based on the equivariant MACE architecture and informed by both ab-initio and semi-empirical data. The resulting potentials reproduce key conformational features of the ApA system, including base stacking, sugar puckering, and backbone flexibility, and provide broader coverage of structural transitions than the general-purpose SO3LR and MACE-OFF24 models. These findings highlight the importance of quantum-accurate RNA force fields towards the structural and energetic characterization of RNA complexes.

## 1. Introduction

The growing understanding of RNA’s role in regulatory and catalytic functions has driven the development of numerous applications in molecular biology and medicine. Notable examples include the advent of mRNA vaccines^1,2^, the development of siRNA therapies^3,4^, and the design of aptamers to target specific molecules^5–10^. These advances emphasize the critical importance of elucidating three-dimensional RNA structure and understanding its interactions with other biomolecules such as lipids and proteins. However, the increasing biological and pharmacological significance of RNA contrasts with the computational resources currently available for its study. As of early 2026, the number of RNA-containing structures in the Protein Data Bank accounted for less than 5%^11^. Given the abundance of canonical double helices in their constitution, there is a much smaller pool of non-trivial motifs, such as junctions, hairpins, and internal loops, available for structural analysis, despite their usual structural and functional relevance^12^. This scarcity represents a major drawback in the development of structure prediction methods, where RNA remains in a template-based stage, and continues to exhibit substantial gaps in the accurate modeling of non-canonical base pairs and non-standard backbone conformations, as concluded by comparative assessments such as CASP and RNA-Puzzles^13,14^. Moreover, it has been reported that the limited availability of experimentally resolved structures is a major hindrance to the training of ML methods for structure prediction^15^.

AAMD, often regarded as the gold standard for exploring biomolecular structure and dynamics, also bears several inherent limitations. Widely used additive atomistic MD force fields, which represent interactions with fixed atomic charges and pairwise additive functional forms, have well-documented deficiencies in describing RNA conformational changes, primarily due to the absence of explicit electronic polarization or many-body interactions^16^. Standard non-polarizable force fields neglect polarization and charge redistribution by construction and instead rely on effective parameter tuning, which cannot fully capture non-additive contributions that arise when the electronic environment changes with conformation or local interactions^17^. This deficiency is particularly important for RNA, as the balance between base stacking, backbone flexibility, ion coordination, and solvation strongly influences the conformational free-energy landscape. Although some improvements have been made in the last few decades to stabilize double helices^18,19^, more complex yet small motifs, such as tetraloops, tetranucleotides, or kink turns, have not been adequately described within a single force-field parametrization^16,20,21^. In particular, imbalances in the relative strength of hydrogen bonds among different molecular components frequently lead to inaccuracies, which are often exacerbated by the choice of water model^22^. For example, conventional pairwise force fields struggle to describe the dynamics of the ribose 2’-hydroxyl and backbone torsions in RNA, both of which are central to correct structural sampling^17^. Neglecting polarization also limits the accurate treatment of divalent cation interactions and charge redistribution, which are critical to RNA folding and stability.

These shortcomings have recently motivated the development of more physically grounded models, including polarizable force fields and ML potentials trained on QM reference data^23–33^. Such approaches can incorporate quantum many-body effects and achieve improved energetic fidelity compared to additive empirical models. In particular, physically inspired ML potentials have shown considerable promise. The general-purpose (atomistic) MACE-OFF24 model^24^, trained on high-fidelity density-functional theory (DFT) data for single drug-like molecules, solvated systems, and molecular dimers, has been demonstrated to accurately describe the dynamical and vibrational properties of large proteins in both gas and condensed phases. Following a similar line of reasoning, the SO3LR model^25^ was recently developed using the SO3krates neural network architecture^34^. By explicitly incorporating both short- and long-range physical interactions and including charged molecules in its training set, SO3LR enabled nanosecond-long DFT-accurate simulations of relevant biomolecules such as the crambin protein, an N-linked glycoprotein, and a lipid bilayer. Similarly, the recently introduced AIMNet2 models^33^ include long-range electrostatics and dispersion interactions in their design. This allows the investigation of organic compounds with diverse charges and valencies, enabling reliable biomolecular simulations of chemically heterogeneous biosystems. Alternatively, Δ-learning approaches that bridge semi-empirical and DFT methods are also emerging as promising strategies to overcome the limitations of classical force fields in biomolecular simulations^35–38^. Despite these advances in physically inspired atomistic ML models, important design challenges remain in ensuring robust sampling of previously unseen conformational changes, such as those commonly encountered in RNA simulations.

To address this challenge, we investigated the impact of QM property data on the performance of ML models for sampling conformational transitions in RNA dimers. The conformations used to develop the ML models were obtained from TREMD simulations of the ApA system. This RNA dimer consists of two nucleosides connected by a phosphate group and represents a simple system that nevertheless exhibits considerable structural richness due to the flexibility of the RNA backbone. Although conformational sampling can be achieved using structure-modeling approaches^39^, we explicitly included solvent molecules to assess their influence on RNA conformations, given the critical role of solvation in mediating RNA interactions^40^. To examine the effect of quantum-level descriptions of interatomic interactions, two different QM methods were used to generate datasets for training the ML models, *i*.*e*., RNA-TB and RNA-DFT (TB stands for tight-binding). The equivariant neural network architecture MACE was employed to train the ML models, and their performance was compared with that of general-purpose ML models, including MACE-OFF24 and SO3LR. Both TREMD and ML-driven simulations revealed six distinct conformational clusters; however, the relative populations of these clusters depend on the ML model used for the MD simulations. Notably, the ML models developed in this work provide broader coverage of the expected structural transitions in the ApA dimer compared to SO3LR and MACE-OFF24. Overall, we expect that this study will motivate the development of computational frameworks for generating comprehensive QM datasets of biomolecular building blocks, thereby enabling more reliable general-purpose ML models for investigating complex RNA systems with unknown structural transitions.

## 2. Methods

### 2.1 Dataset Generation

#### Conformational sampling strategy

Following the procedure reported by Hayatshahi *et al*.^41^, we performed TREMD simulations^42,43^ to generate an extensive sample of the conformational space of the ApA dimer. In TREMD, multiple replicas of the system are simultaneously simulated at different temperatures. At fixed intervals of time, a temperature exchange between two randomly chosen, adjacent replicas is attempted and accepted according to a Metropolis-like criterion that depends on the energy and temperature of the replicas. Denoting the temperatures by *T*_*i*_ and *T*_*j*_, and the potential energies by *U*_*i*_ and *U*_*j*_ of replicas *i* and *j* respectively, the probability of accepting an exchange is given by^42,43^,

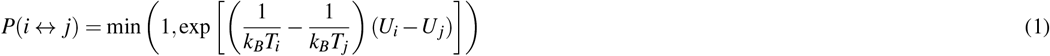

where *k*_*B*_ is Boltzmann’s constant. This protocol allows each replica to traverse between high and low temperatures, thereby enhancing sampling efficiency. The high-temperature replicas can overcome energy barriers, while low-temperature replicas, usually close to room or physiological conditions, provide a finer sampling of the local conformational landscape. For subsequent analysis, only the configurations of the replicas belonging to the room temperature are considered.

In our setup, 18 independent simulations were run for 500 ns over a wide range of temperatures, spanning a range between 280 K and 396.4 K with a geometric distribution (280, 285.8, 291.7, 297.7, 303.9, 310.1, 316.6, 323.1, 329.8, 336.6, 343.5, 350.6, 357.9, 365.3, 372.8, 380.5, 388.4 and 396.4 K). Room temperature was chosen to be 297.7 K and temperature exchange between replicas was attempted every 500 integration steps. The equations of motion were integrated using the leapfrog method with a time step of 2 fs, saving the coordinates every 5000 integration steps. In all cases, the temperature was controlled using the Stochastic Velocity-Rescaling thermostat^44^ with a time constant of 0.1 ps. The simulations were performed in the GROMACS 2023^45^ simulation package using the TIP3P water model^46^ with the AMBER99 force field^47^, using parmbsc0^18^ and *χ*OL3 corrections^48^. The diagonal terms of the empirical transition matrix exhibited values between 0.88 and 0.67, with an average of 0.73. Before the TREMD simulations, the system started from an A-form configuration, whose energy was minimized in vacuum by steepest descent for 5000 steps. After the introduction of 1080 water molecules and a single Na ion^49^ to neutralize the system charge, the energy was minimized once again with the same method for 4,000 steps, followed by a short equilibration run at constant temperature and pressure, using the Stochastic Velocity-Rescale thermostat (time constant of 0.1 ps and temperature of 300 K) and Berendsen barostat (target pressure of 1 bar and time constant of 1 ps)^50^. After this, the molecules were enclosed in a cubic simulation box with a side of 32.1719 Å.

To validate the conformational space sampled by TREMD simulations, an unbiased MD simulation (plain-MD, PMD) was performed as an NVT simulation at a temperature of 297.7 K, using the same thermostat as in the TREMD simulations and following the same equilibration procedure. The production run was carried out for a total duration of 4 *µ*s, which is a time scale shorter than the necessary time for achieving full conformational convergence, as reported by previous studies^41^. The purpose of this choice is to evaluate the impact of using an incomplete sampling of the conformational space on the structural features and the training of the force field.

#### Quantum-mechanical calculations

For the next step in the dataset generation, we have considered only the TREMD trajectory produced at 297.7 K. From the whole trajectory, we have selected conformations saved every 10 ps, making a total of 50,000 initial conformations. These conformations were later classified into six clusters, representing the possible structural conformations adopted by the ApA dimer (*vide infra*). We then extracted the ApA dimer and a water shell of 3 Å around it from these conformations using the VMD package. To refine this extensive set of conformations, we applied an RMSD-based hierarchical clustering method. This step produced 35,348 structures, ensuring that the chosen structures effectively represent the diversity of the explored conformational space. The number of selected structures per conformational cluster (*vide infra*) are, Aform:3,216, Inverted: 10,997, Ladder: 4,958, Anti-ladder: 2,240, Sheared: 3,793, and Unstacked: 10,144.

Another goal of our work is to examine the best routes for generating accurate QM datasets of RNA systems. In this context, we have generated two QM datasets using the semi-empirical density functional tight-binding (DFTB) method^51^ and the more accurate density functional theory (DFT). Accordingly, energies, atomic forces, and atomic charges of solvated ApA conformations were calculated using the third-order DFTB method (DFTB3)^52^ supplemented with a treatment of many-body dispersion (MBD) for van der Waals interactions^53,54^, as it is implemented in the DFTB+ code^55^. Single-point DFTB3+MBD calculations were carried out by considering hydrogen correction and the electronic Hamiltonian described by the 3ob parameters set^56^. Similar to previously developed datasets of (bio)molecular systems^27,57,58^, single-point DFT calculations were computed using the hybrid PBE0 functional^59,60^ supplemented with MBD correction using the `FHI-aims` code^61,62^ (version 180218). For these calculations, “tight” settings were applied for basis functions and integration grids. For both types of calculations, we have excluded the Na ion from the calculations; *i*.*e*., the ApA dimer together with the water shell was considered as a charged system with net charge *Q* = −1.0. Atomization energies for both QM datasets were obtained by subtracting the atomic DFTB3 and PBE0 energies from the corresponding total energy of each molecular conformation.

### 2.2 Clustering technique

To refine the description of the ApA conformations, the analysis was restricted to nucleobase conformations, and stacking interactions were identified using the Dissecting the Spatial Structure of RNA (DSSR) approach^63^. This decision enabled a more precise structural characterization than standard global clustering methods. This screening process yielded 3,132,000 stacked frames from the 4 *µ*s PMD simulation, saved every 1 ps, and 39,193 frames from the TREMD ensemble. Using the CPPTRAJ module of AmberTools24, the remaining frames were aligned by centering both residues and utilizing a specific subset of heavy atoms from the nitrogenous bases (C2, C4, C5, C6, C8, N1, N3, N6, N7, and N9) (see Table S1 of the Supplementary Information (SI)). This selection was specifically designed to emphasize the relative orientation and stacking geometry while minimizing the influence of the flexible sugar-phosphate backbone.

Pairwise RMSD values were calculated across the aligned frames, and the conformational space was partitioned into seven clusters using the k-means algorithm with random centroid initialization and complete linkage. Fixing the number of clusters at *k* = 7 across all systems ensured statistical consistency for subsequent comparisons. These seven clusters were eventually grouped into six primary conformational families determined by the mutual orientation of the nucleobases. This approach introduces key modifications to the framework established by Hayatshahi et al.^41^.

While our study utilized k-means clustering, Hayatshahi et al. employed a different methodology and atom set defined by C5^′^, C4^′^, C3^′^, O3^′^, P, and C2 that correspond to a selection of positions in the RNA backbone and the nitrogenous base (see Table S1 of the SI) and adaptively tuned the number of clusters between four and seven for each system. In contrast, our methodology utilizes a rigid cluster count and a base-centric atom selection to focus specifically on *π*-stacking interactions. Despite these procedural differences, the resulting centroids displayed significant structural similarity to those previously reported. Consequently, the nomenclature from the previous literature was adopted to describe the five final groups: A-form, inverted, ladder, sheared, and anti-ladder.

### 2.3 Machine learning potentials

We employed the state-of-the-art equivariant neural network MACE^64^ to develop ML potentials for exploring the potential energy surface of the ApA dimer. MACE builds upon the mathematical framework of the Atomic Cluster Expansion (ACE) and incorporates multiplicative interactions between geometric and atomic features. To assess how the level of electronic-structure theory influences the prediction of structural transitions in the ApA dimer, separate ML potentials were trained on datasets generated using DFTB and DFT. Each dataset was divided into 80%, 10%, and 10% to generate the training, validation, and test sets, respectively. Key hyperparameters, including the cutoff radius (*r*_c_), *l*_max_, and the number of interaction layers (*N*_int_), were optimized to identify the best-performing ML potential for each QM dataset. The number of feature channels was fixed at 128 for all models.

After identifying the best-performing models, we performed constant-temperature MD simulations of the gas-phase ApA dimer. As the initial structure, we selected a conformation from the A-form cluster, the only structure that has been experimentally resolved^65^. For each model, three independent MD simulations were carried out at 300 K for 10 ns using a Langevin thermostat as implemented in the ASE package^66^. Different initial velocity distributions were used in each run. The timestep was set to 0.5 fs, and the friction coefficient was 2 × 10^−3^. To improve the sampling of structural transitions in the ApA dimer, the three trajectories were concatenated, yielding a total simulation time of 30 ns per ML model. To validate the performance of the RNA-TB and RNA-DFT models, we also performed MD simulations using two recently developed general-purpose atomistic ML potentials for biomolecular systems: MACE-OFF24^24^ and SO3LR^25^. Since MACE-OFF24 was trained exclusively on neutral molecules and does not include Na atoms, the phosphate group was protonated to neutralize the ApA dimer. In contrast, SO3LR supports charged systems, so the initial structure with total charge *Q* = −1e was used without modification. All geometries were optimized with the corresponding method prior to the MD simulations.

### 2.4 Hellinger analysis of distributions

The Hellinger distance was used to quantify the similarity between dihedral angle distributions obtained for each RNA model relative to the TREMD simulation (reference). This metric is a well-defined statistical measure for assessing the similarity between two probability distributions^67^.

For discretely sampled MD data, the Hellinger distance between two normalized distributions *p* = {*p*_*i*_} and *q* = {*q*_*i*_} is computed as

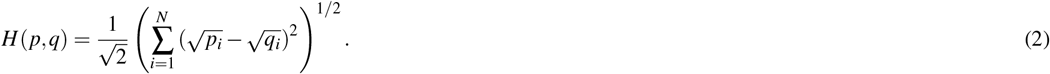

Here, *p*_*i*_ and *q*_*i*_ denote the normalized histogram probabilities of the two dihedral angle distributions. The metric satisfies 0 ≤ *H* ≤ 1, where *H* = 0 indicates identical distributions and *H* = 1 corresponds to distributions with no overlap.

The Hellinger distance quantifies the overlap between fluctuation profiles and thus provides a robust metric for comparing two simulations with respect to their sampled conformational space. Empirically, values below 0.3 indicate excellent agreement, whereas values up to approximately 0.7 still reflect acceptable similarity.

## 3 Results & Discussion

### 3.1 Cluster characterization

Fig. 1(b) summarizes definitions of the clusters and the conformation populations sampled by the ApA dimer during simulations performed with PMD and TREMD. As explained in the Methods section, we adopt a similar nomenclature to the one proposed by Hayatshahi *et al*.^41^, identifying A-form (A), sheared (S), inverted (I), ladder (L) and unstacked (U) clusters. In addition, we observed a small cluster that we denominate “anti-ladder” (AL) since its centroid is characterized by a different orientation of the nucleobases with respect to the ladder cluster. The most frequently visited conformation in both protocols was *inverted*, accounting for 36 % of frames in PMD and 30 % in TREMD. The A-form conformation was observed in 12 % of frames under both conditions, indicating a consistent presence regardless of the sampling strategy. Ladder conformation was moderately observed at 13 % and 8 % in PMD and TREMD, respectively. A similar trend was seen for the *sheared* conformation, which decreased from 10 % in PMD to 6 % in TREMD. The anti-ladder conformation appeared at equal frequencies (6 %) in both methods. Notably, unstacked conformations were more prevalent in TREMD (39 %) than in PMD (22 %), suggesting that the broader conformational exploration by TREMD enhances the sampling of unstacked conformations. The results indicate a similar qualitative ensemble, but replica exchange tends to shift the population balance towards the unstacked conformation.

**Figure 1.**
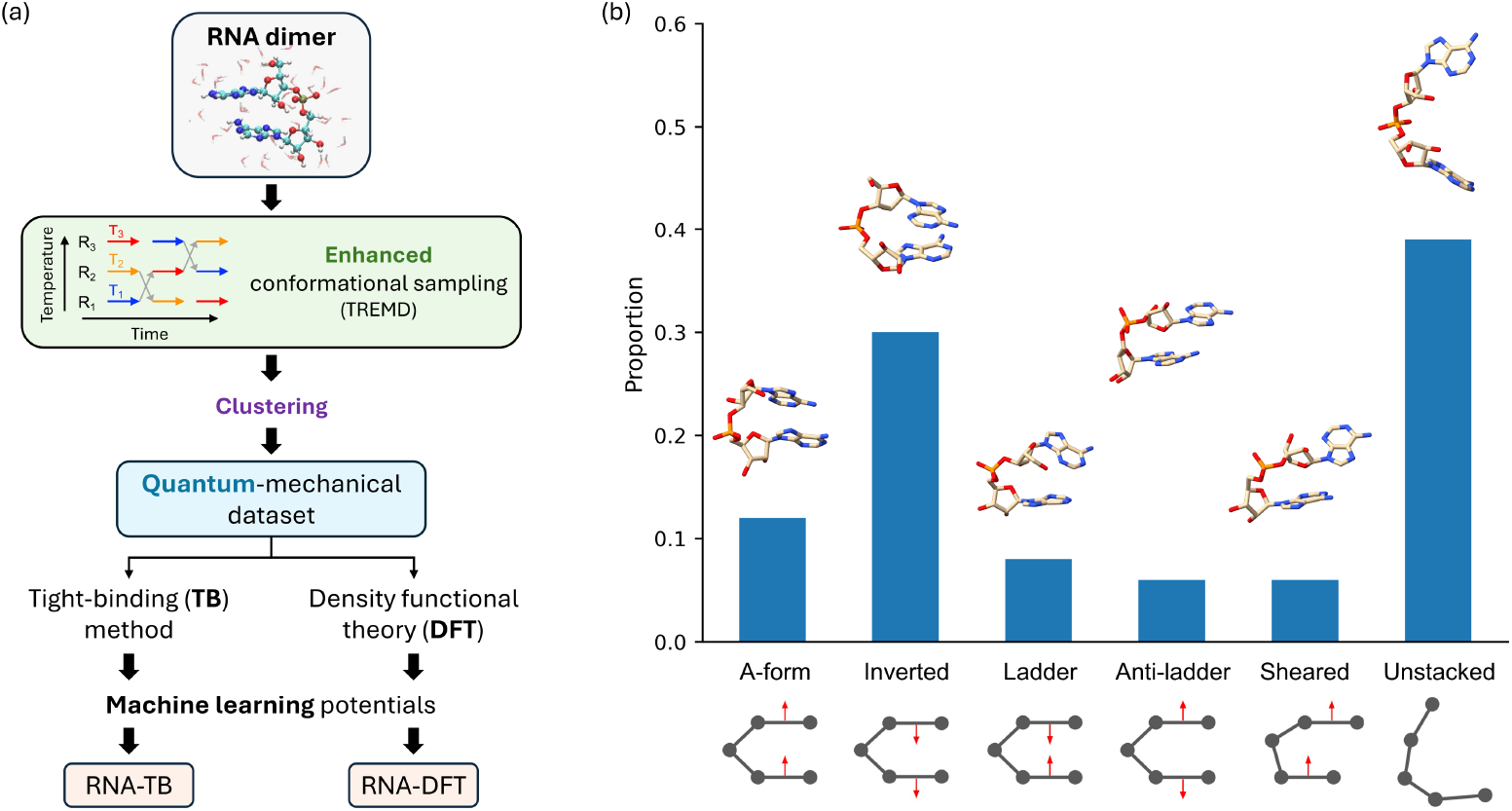
(a) Workflow for the development of ML potentials for RNA modeling. The models were trained on a QM dataset generated using two electronic-structure methods: density functional tight-binding (DFTB) and density functional theory (DFT). (b) Conformational classification and populations of ApA obtained from TREMD simulations. Six conformational clusters were identified using the procedure described in Sec. 2.2: A-form (A), inverted (I), ladder (L), anti-ladder (AL), sheared (S), and unstacked (U). Representative structures of each cluster are shown.

We next analyze the QM-derived properties of the conformations within each cluster. Fig. 2(a) presents boxplots of the atomization energy, *E*_AT_, for solvated ApA dimers computed using PBE0+MBD (DFT) and DFTB3+MBD (TB). As expected, the two electronic-structure methods yield clearly distinct energy distributions, with mean values of ⟨*E*_AT_⟩ =-1,036.6 eV for DFT and -761.3 eV for TB. This systematic offset primarily reflects differences in the underlying electronic Hamiltonians, including the all-electron treatment and exchange–correlation description in PBE0 compared to the parameterized, approximate framework of DFTB3. The spread of the energy distributions also differs between methods. DFT energies exhibit a standard deviation of *σ*_E_ = 52.2 eV, whereas the corresponding value for TB data is reduced to 39.2 eV. This larger variance in the DFT results indicates a stronger sensitivity to conformational changes, likely arising from a more detailed treatment of electronic structure and polarization effects. Focusing on the van der Waals (vdW) contribution, DFTB3+MBD yields slightly smaller mean MBD energies than PBE0+MBD, with *E*_*MBD*_⟩ = -5.06 eV and -5.62 eV, respectively. These differences originate from the distinct approximations used to compute atomic charges in each method, which directly enter the MBD formalism and influence the resulting MBD energy. In contrast, Lennard–Jones (LJ) energies are positive and lie well outside the energy range predicted by both QM approaches, underscoring the fundamentally different treatment of long-range dispersion interactions in the AMBER99 force field, where dispersion is modeled via pairwise additive terms rather than an explicit many-body formalism.

**Figure 2.**
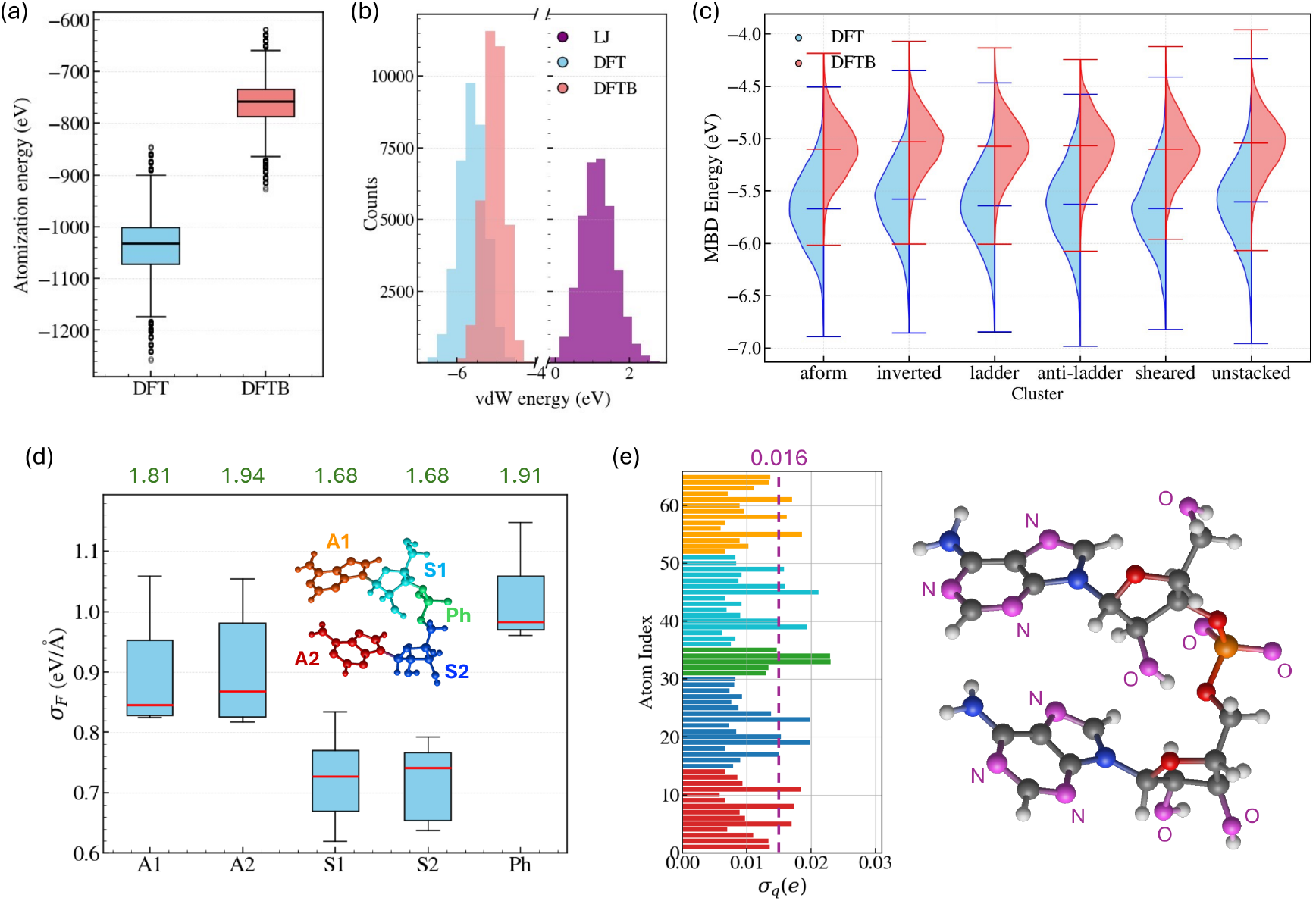
(a) Box plots of the atomization energies computed with DFT (PBE0+MBD) and DFTB (DFTB3+MBD) for ApA conformations sampled from the reference TREMD simulation. (b) Frequency distributions of the van der Waals (vdW) energies for ApA conformations computed with DFT, DFTB, and the Lennard-Jones potential of the AMBER99 force field. (c) Distributions of the MBD energies by conformational cluster obtained with DFT and DFTB. (d) Box plots of the standard deviation of atomic forces, *σ*_F_, computed with DFT for the ApA building blocks: adenine 1 (A1), adenine 2 (A2), sugar 1 (S1), sugar 2 (S2), and phosphate (Ph). Mean force values for each group are also shown on the top of box plots. (e) We show the variance of atomic charges, *σ*_q_, computed with DFT for all atoms in ApA dimer. The atom index has been ordered with respect to the molecular groups defined in panel (d). In the inserted structure, we have highlighted in violet the atoms with *σ*_q_ > 0.016e.

A cluster-resolved analysis of the MBD energies further reveals that discrepancies between DFTB3+MBD and PBE0+MBD become more pronounced for structurally complex systems, particularly stacked and unstacked conformations of the solvated (negatively charged) ApA dimer (see Fig. 2(c)). Notably, the MBD correction within DFTB3 was originally parameterized to reproduce interaction energies of S66x8 molecular dimers computed at the PBE0+MBD level^55^, suggesting that deviations may increase in larger or more heterogeneous environments. Similar discrepancies between PBE0+MBD, DFTB3+MBD, and the AMBER force field have recently been reported for ligand–pocket interaction energies^68^. Finally, an analysis restricted to conformations containing 194 atoms shows that structures belonging to the A-form and Inverted clusters exhibit the largest MBD energies among all clusters (see Fig. S1 of the SI), consistent with their more compact and strongly stacked geometries.

Fig. 2(d) shows the fluctuations of the standard deviation of the atomic forces, *σ*_F_, for the five building blocks of the ApA dimer: Adenine 1 (A1), Adenine 2 (A2), Sugar 1 (S1), Sugar 2 (S2), and Phosphate (Ph) groups. Following our previous analysis, we report only the results obtained at the PBE0+MBD level (see DFTB results in Fig. S2 of the SI). Although the DFT calculations include explicit water molecules, the analysis focuses exclusively on the ApA atoms. We find that atoms in the Ph group exhibit the largest force fluctuations (high *σ*_F_), together with a high mean force of ⟨*F*⟩ = 1.91 eV/Å. This behavior likely reflects the central role of the Ph group in the structural transitions between conformational clusters sampled during the TREMD simulation. In addition, the adenine groups display larger ⟨*F*⟩ and *σ*_F_ values compared to the sugar groups. A similar trend is observed in the TB data, suggesting that it originates from the strong and multiple interactions between adenine atoms and the surrounding water molecules. Regarding atomic charges, *q*, computed at the PBE0+MBD level, most atoms in the ApA dimer exhibit a substantial variance in charge magnitude, *σ*_q_ (see Fig. 2(e)). In particular, two O atoms in the Ph group show the largest *σ*_q_ values (green bars in the left panel of Fig. 2(e)), which can be associated with the overall negative charge of the system. Furthermore, the O atoms in the hydroxyl groups of the sugars and the N atoms in the adenine groups also display large charge fluctuations, likely due to strong interactions with surrounding water molecules (see violet balls in the inset of Fig. 2(e)). In contrast to the DFT results, the TB data do not show large *σ*_q_ values for the O atoms in the sugar groups relative to atoms in the adenine and phosphate groups (see Fig. S2 of the SI). These differences are expected to impact the performance of ML potentials trained on DFT versus TB data. Moreover, since classical force fields employ fixed atomic charges during MD simulations, our results highlight the importance of incorporating QM effects into force-field development to accurately describe interatomic interactions in RNA structures.

### 3.2 Predicting the potential energy surface

We now examine how different electronic structure references influence the performance of the ML potentials in sampling the conformational space of the ApA dimer. To parameterize these potentials, all conformational clusters were merged into a single dataset, which was then randomly divided into training, validation, and test subsets. Following an extensive hyperparameter optimization, we report results obtained with a fixed number of interactions (*N*_int_ = 2) and maximum angular momentum (*l*_max_ = 1), while varying the cutoff radius (*r*_c_) between 4.0 and 6.0 Å. The mean absolute errors (MAEs) for ML potentials trained on the TB- and DFT-based datasets are summarized in Table 1. In both cases, the MAEs for energies and forces decrease as *r*_c_ increases. A particularly pronounced improvement is observed when increasing *r*_c_ from 4.0 to 5.0 Å, indicating the importance of including longer interatomic interactions. The most accurate models are obtained with *r*_c_ = 6.0 Å, yielding energy MAEs of 1.58 and 2.10 mkcal/mol per atom for the RNA-TB and RNA-DFT models, respectively. The force predictions are similarly accurate, with MAEs of 0.087 and 0.108 kcal/mol Å, respectively.

**Table 1.** Evaluation of the performance of ML potentials trained on DFTB3+MBD (RNA-TB) and PBE0+MBD (RNA-DFT) property data for the ApA system. The equivariant MACE architecture was employed for model development, and the cutoff radius (*r*_c_) was optimized. The number of interaction layers was set to *N*_int_ = 2, with *l*_max_ = 1. Mean absolute errors (MAEs) in the prediction of total energies and atomic forces are reported for all ApA conformations, as well as MAEs in atomic forces resolved by conformational cluster. For comparison, results obtained with the general-purpose SO3LR model are also shown. Errors for energies and atomic forces are reported in mkcal/mol per atom and kcal/mol·Å, respectively.

| Model | Hyperparameter | | | Full | | error in $F_{\text{at}}$ per cluster | | | | | | |
| --- | --- | --- | --- | --- | --- | --- | --- | --- | --- | --- | --- | --- |
| | $N_{\text{int}}$ | $l_{\text{max}}$ | $r_c$ | $E$ | $F_{\text{at}}$ | A | I | L | AL | S | U | |
| RNA-TB | 2 | 1 | 4.0 | 8.33 | 0.329 | 0.326 | 0.322 | 0.326 | 0.331 | 0.327 | 0.339 |  |
|  | 2 | 1 | 5.0 | 2.83 | 0.138 | 0.135 | 0.133 | 0.136 | 0.138 | 0.137 | 0.145 |  |
|  | 2 | 1 | 6.0 | **1.58** | **0.087** | 0.085 | 0.083 | 0.085 | 0.087 | 0.086 | 0.093 |  |
| RNA-DFT | 2 | 1 | 4.0 | 9.64 | 0.321 | 0.319 | 0.315 | 0.318 | 0.323 | 0.320 | 0.328 |  |
|  | 2 | 1 | 5.0 | 3.59 | 0.153 | 0.151 | 0.149 | 0.151 | 0.154 | 0.152 | 0.159 |  |
|  | 2 | 1 | 6.0 | **2.10** | **0.108** | 0.107 | 0.105 | 0.106 | 0.109 | 0.108 | 0.112 |  |
| SO3LR | - | - | - | - | 1.836 | 1.863 | 1.880 | 1.828 | 1.830 | 1.816 | 1.793 |  |

By analyzing the performance across conformational clusters, we find that stacking conformations are better described by the ML potentials than unstacked ones. This trend likely arises from the more compact nature of stacking conformations, which feature a higher number of short interatomic distances. In contrast, the reduced accuracy observed for unstacked conformations highlights limitations of the ML models in describing long-range interatomic interactions. For comparison, we also evaluate the general-purpose SO3LR model on the DFT dataset, as it was trained at the PBE0+MBD level of theory and is applicable to charged systems such as the ApA dimer. Our analysis focuses on force predictions, since SO3LR was trained exclusively on atomic forces from approximately four million equilibrium and non-equilibrium molecular structures. SO3LR yields an MAE of 1.836 kcal/mol·Å, which is higher than that obtained with our ML models but still moderate. This discrepancy may originate from the complex interatomic interactions surrounding the negatively charged phosphate group, which is underrepresented in the dataset used to parameterize SO3LR. In contrast to the RNA-TB and RNA-DFT models, however, SO3LR provides more accurate predictions for conformations in the unstacked cluster, likely reflecting its improved treatment of long-range interatomic interactions by design.

### 3.3 Conformational transitions between ApA clusters

To evaluate how accurately the developed ML potentials describe the conformational landscape of the ApA dimer, we performed gas-phase MD simulations at 300 K and analyzed the stacking fraction *F*_s_ as well as structural transitions. All simulations were initiated from the stacked A-form configuration (see Sec. 2.3 for computational details). To quantify conformational transitions, each frame of the resulting MD trajectories was analyzed using the Dissecting the Spatial Structure of RNA (DSSR) method^63^. Each snapshot was systematically classified as either stacked or unstacked based on the geometric orientation of the nucleobases. Specifically, DSSR identifies stacking by calculating the vertical distance and the horizontal displacement between adjacent base planes, ensuring that the classification reflects a physical overlap of the aromatic rings rather than mere proximity. For frames identified as stacked, we further categorized the specific geometry by aligning the heavy atoms of the nitrogenous bases (C2, C4, C5, C6, C8, N1, N3, N6, N7, and N9) to five reference configurations: A-form (A), inverted (I), ladder (L), sheared (S), and anti-ladder (AL). RMSD was calculated for each frame against these five templates using the common atom set. This process generated five distinct RMSD time series for every trajectory. Each frame was then assigned a conformational label. Snapshots were passed by DSSR and results with no stacking were classified as unstacked, while all other frames were assigned the label of the reference geometry that yielded the lowest RMSD value. This classification enabled a systematic comparison of stacking populations, unstacked lifetimes, and transition probabilities across all models.

Fig. 3(a) compares the concentration of A-form conformations (*C*_A_) and the stacking fraction (*F*_s_) obtained using the RNA-TB and RNA-DFT models. The A-form conformation was selected because it is the only structure that has been experimentally resolved^65^. We also include results from the reference TREMD simulation and the general-purpose ML models MACE-OFF24 and SO3LR. The symbol size represents the MAE in force predictions. Among the RNA-TB models, the model trained with a cutoff radius of *r*_c_ = 5.0 Å yields *F*_s_ and *C*_A_ values closest to those obtained from the TREMD simulation (*F*_s_ = 0.78 and *C*_A_ = 12%, shown as pink dashed lines). For the RNA-DFT models, the best agreement with the reference data is achieved for *r*_c_ = 6.0 Å. Because these two models reproduce both *F*_s_ and *C*_A_ most accurately, we use them for the remaining analyses. SO3LR exhibits the largest stacking fraction among all ML models; however, it shows a strong preference for the A-form, with *C*_A_ = 54%, which is not observed in the reference results. In contrast, despite predicting a large stacking fraction, MACE-OFF24 fails to reproduce the A-form population, yielding *C*_A_ = 1%. Based on these observations, MACE-OFF24 is excluded from further analysis.

**Figure 3.**
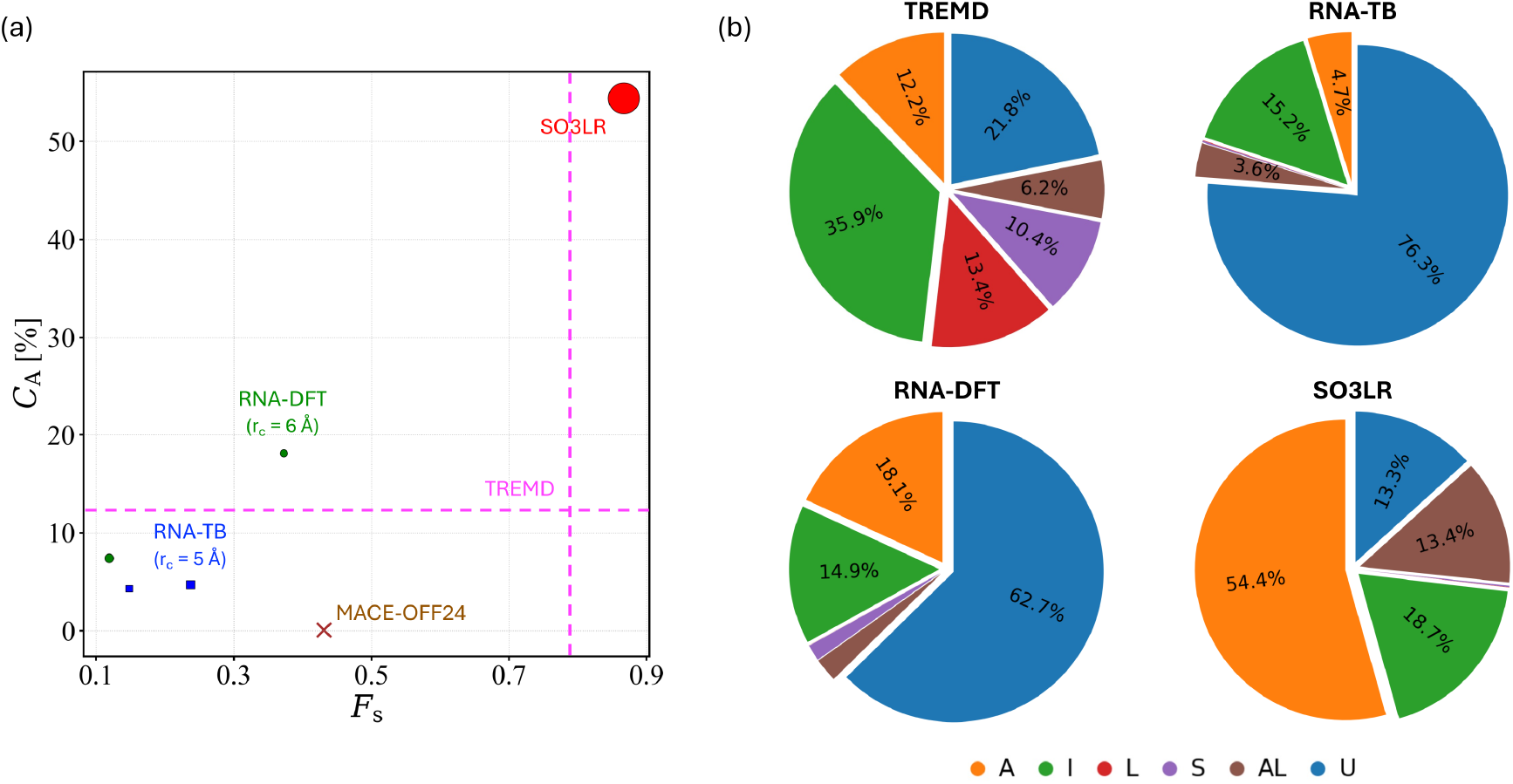
(a) Comparison between the concentration of A-form conformations (*C*_A_) and the stacking fraction (*F*_s_) obtained from MD simulations using the RNA-TB, RNA-DFT, and general-purpose ML models (MACE-OFF24 and SO3LR). Results are shown for two cutoff radii (*r*_c_): 5.0 Å and 6.0 Å. Symbol sizes scale with the mean absolute error (MAE) in atomic force predictions for all ApA conformations (see Table 1), except for the MACE-OFF24 model. Results from the reference TREMD simulation are indicated by pink dashed lines. (b) Pie charts showing the relative populations of the six conformational clusters obtained from the corresponding MD simulations: A-form (A), inverted (I), ladder (L), anti-ladder (AL), sheared (S), and unstacked (U). Results from the reference TREMD simulation are included for comparison.

We now examine the conformational preferences for the stacking geometries of ApA dimers obtained by the selected ML models and the reference TREMD simulation, see Fig. 3(b). In TREMD, the conformational ensemble is heterogeneous, dominated by inverted (36 %) and unstacked (22 %) states, with moderate populations of A-form (12%), sheared (10%), and ladder (13%) conformations. In contrast, the SO3LR model favors A-form stacking (54%) and includes moderate populations of inverted (19%), anti-ladder (13%), and unstacked (13%) conformations but does not sample ladder or sheared conformations. The RNA-TB model displays a strong preference for unstacked geometries (76%), followed by minor populations of inverted (15%) and anti-ladder (4%) conformations, and virtually no sampling of A-form or ladder states. The RNA-DFT model also favors unstacked configurations (63%), with inverted conformations as the second most populated conformation (15%). Also, this model samples A-form (18%), anti-ladder, ladder, and sheared states. Note that TREMD exhibits a more heterogeneous ensemble where the inverted conformation is the most prevalent at 36%, supporting the presence of multiple transient stacked states rather than a single dominant geometry. However, RNA-TB and RNA-DFT models show that the unstacked conformation dominates the population, indicating that these models likely underestimate the strength of base-stacking interactions or fail to capture the stability of the stacked conformations.

Fig. 4 shows the transition matrices derived from three concatenated 10-ns simulations per model, sampled at 10-fs intervals. The study of ApA dimers provides a simplified yet biophysically rich model for exploring the elementary processes of base stacking and unstacking. These fluctuations represent the primary steps towards large-scale conformational rearrangements^69^. The analysis of transition matrices derived from these simulations reveals stark differences in how each model explores the conformational landscape. While TREMD enhances conformational sampling by allowing exchanges between replicas at different temperatures, this facilitates barrier crossing at the expense of temporal consistency. Because these exchanges disrupt the continuous physical timeline, trajectories generated through TREMD are fundamentally unsuitable for direct kinetic analysis. In contrast, the PMD simulation preserves full dynamical continuity. As a classical MD approach, it provides a seamless timeline while remaining sufficiently long to sample the relevant conformational space. This allowed the PMD trajectory to be used for computing the transition matrix, enabling a reliable characterization of the interconversion among the six conformations. This diversity is essential for biological function, where structural flexibility allows RNA to adapt to protein binding or ligand intercalation^70^. In the SO3LR model, a high number of transitions is observed specifically between the A-form and the inverted or anti-ladder conformations. This behavior suggests that in SO3LR, the activation barriers between these conformations are relatively low or the potential energy surface is sufficiently smooth to allow frequent transitions. The transition matrices for RNA-TB and RNA-DFT are characterized by sparse exchanges and frequent movement to and from the unstacked conformation. This suggests a sink where, once the base-stacking interaction is disrupted, the model lacks the directional force to return to a canonical stack. The conformational transitions of RNA dimers are governed by a delicate interplay of quantum mechanical forces that define a rugged and multifaceted energy landscape. The analysis of transition matrices reveals that while TREMD provides a comprehensive view of this landscape, ML potentials like SO3LR are beginning to capture the dynamic essence of stacked-substate interconversion. The ability of SO3LR to sample A-form and anti-ladder transitions suggests a high-fidelity representation of the glycosidic torsion and many-body dispersion interactions, even if its exploration remains localized. In contrast, the bias of RNA-TB and RNA-DFT toward unstacked conformations highlights the challenges of modeling RNA dynamics without explicit solvation or advanced many-body terms. The sparse transitions in these models reflect a landscape where the pathways between canonical and non-canonical geometries are obstructed by overestimated barriers or missing solvent-driven stabilizing forces.

**Figure 4.**
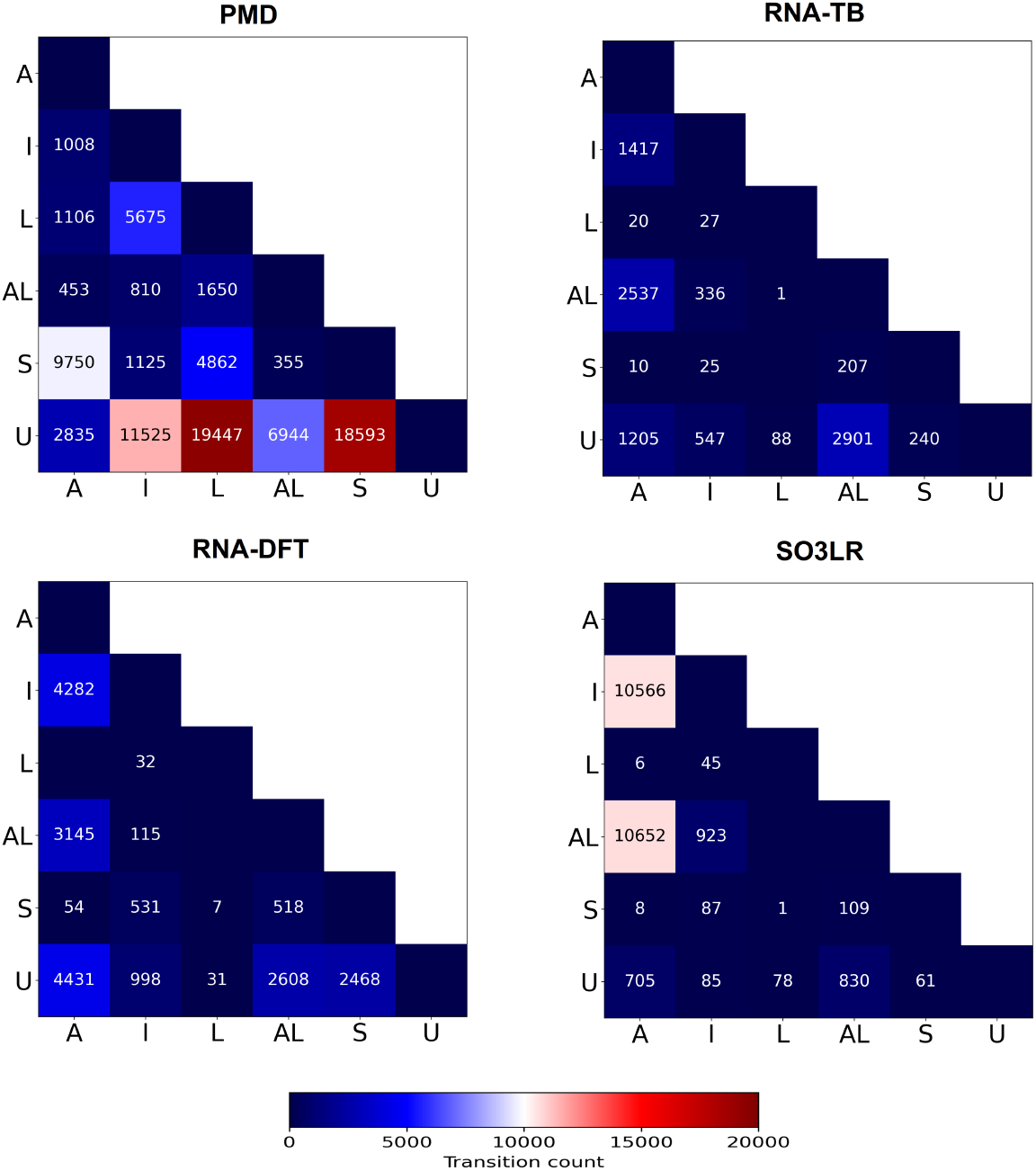
The transition behaviors for ApA dimers are compared across various models. The stacking transition matrices illustrate the frequency of transitions between stacking states for each model. Each panel represents a different simulation method. Data for RNA-TB, RNA-DFT, and SO3LR was accumulated from three 10-ns replicas, sampled every 10 fs to yield 10^6^ frames per model. The PMD model was sampled over a 4-*µ*s trajectory with a 1-ps saving interval, resulting in 4 × 10^6^ frames.

### 3.4 Comparative analysis of RNA dihedrals

To assess the ability of different ML models to capture the conformational preferences of the ApA dimer, we analyzed the distribution of glycosidic torsion angles (*i*.*e*., *χ* and *δ* dihedral angles) for the two nucleotides (see Fig. 5(a)). Thus, we have computed two-dimensional probability maps for these dihedral angles. Fig. 5 shows the distributions obtained from the TREMD simulations, which serve as our reference. Fig. 5 (b) shows the two-dimensional density plots for *χ*_1_ and *χ*_2_ distributions, which are characterized by a dominant population in the anti–anti region (around −60°, −180°)^71^, reflecting the canonical geometry expected for A-form RNA dimers. The distributions indicate that both nucleotides maintain relatively stable and planar orientations. In comparison, the SO3LR model recovers the general location of the dominant basin, but the sampled distribution is shallow and slightly shifted to very planar geometry −180°. Notably, the density extends into less favorable regions of the torsional space, suggesting that SO3LR underestimates conformational flexibility and includes non-native rotamers. The RNA-TB model shows a sharper, more localized distribution centered in the same anti-anti region observed in the TREMD simulations. The close agreement in both location and shape of the distribution suggests that the RNA-TB model effectively captures the native torsional preferences of RNA bases, providing a more efficient sampling of the nucleotide conformations. Finally, the RNA-DFT model performs better along the *χ* angles, sampling the anti–anti region in better agreement with the TREMD reference. These findings highlight potential limitations in the transferability of the torsional parameters associated with the *χ* dihedral angles in the general-purpose SO3LR and MACE-OFF24 models (see Fig. S8).

**Figure 5.**
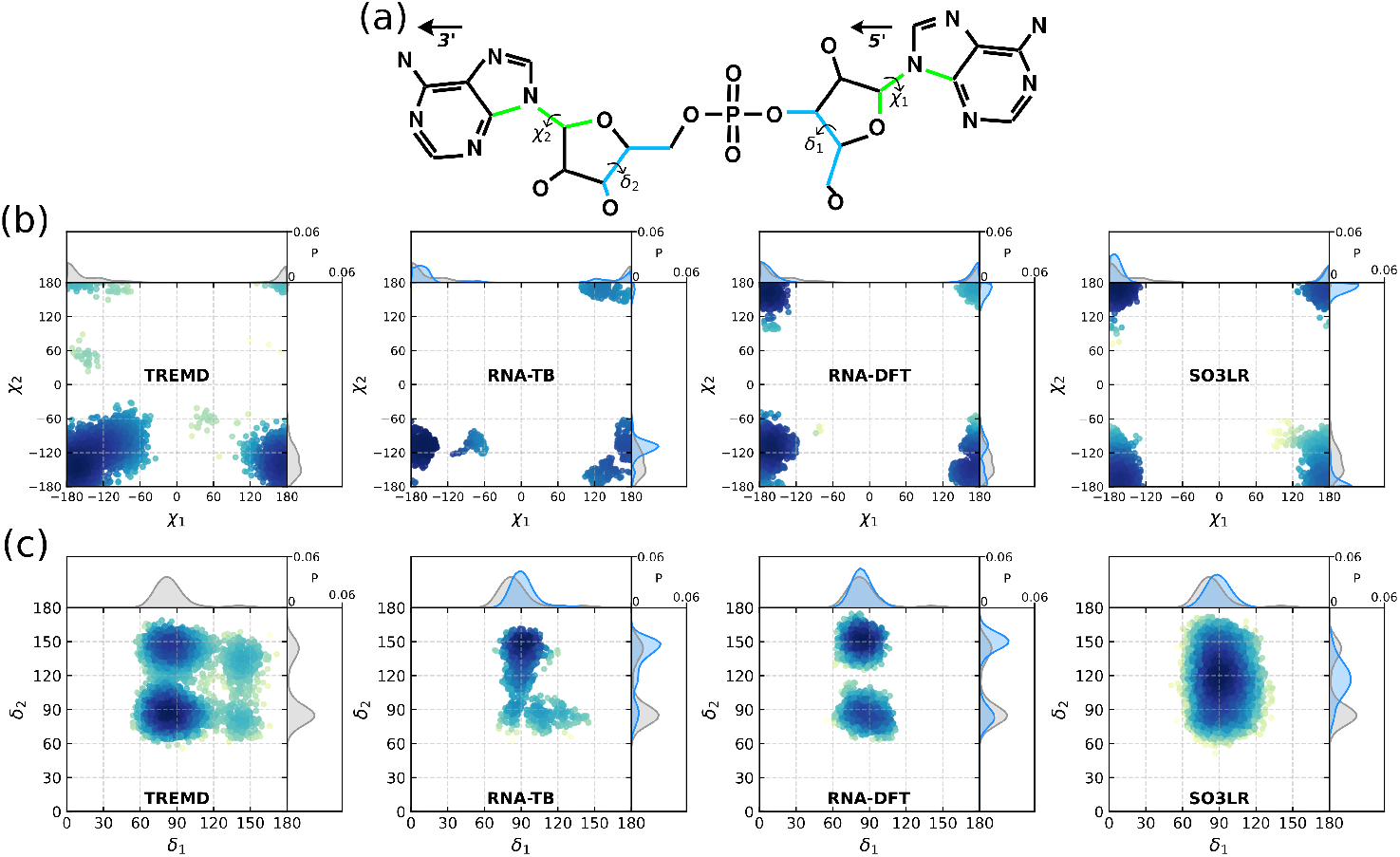
(a) Chemical structure of the ApA RNA dimer showing the definition of the *χ* and *δ* torsion angles for both nucleotides. (b) Two-dimensional distributions of *χ*_1_ and *χ*_2_ angles sampled during simulations using the TREMD, RNA-TB, RNA-DFT, and SO3LR models. (c) Two-dimensional distributions of *δ*_1_ and *δ*_2_ angles for the same models. The color scale represents the normalized probability of sampling each pair of torsional angles.

Fig. 5 (c) displays two-dimensional density plots of *δ*_1_ and *δ*_2_ torsion angles for each nucleotide in the ApA dimer. The TREMD reference data exhibits a pronounced bimodal distribution along *δ*_2_ and slight one along *δ*_1_. A dominant population is centered near *δ*_1_ ≈ 85° and *δ*_2_ ≈ 85°, where the associated *δ*_1_ and *δ*_2_ angles fall within the 55° to 110° range^65,72^. This conformation is characteristic of canonical A-form RNA, where both sugar puckers remain in the lower *δ* range. A secondary population is identified around *δ*_1_ ≈ 85° and *δ*_2_ ≈ 145°, indicating a configuration where *δ*_1_ maintains a value between 55° and 110° while *δ*_2_ adopts a value in the 120° to 175° range^65,72^. This specific arrangement results in an asymmetric conformation that often facilitates transitions between conformations ^65^. The SO3LR model shows a broad distribution centered around *δ*_1_ ≈ 90° and *δ*_2_ ≈ 120°. This indicates that this model preserves the general A-form sugar geometry while allowing more flexibility in puckering compared to the reference TREMD data. RNA-TB presents a distribution centered near *δ*_1_ ≈ 90° and *δ*_2_ 1≈ 50°, consistent with a strong preference for this dihedral angles A minor shoulder was observed around *δ*_1_ ≈ 90° and *δ*_2_ ≈ 85°, which follows the symmetric C3^′^-endo sugar pucker conformation observed in TREMD. The RNA-DFT model closely reproduces the bimodal distributions observed in the TREMD reference, with two distinct populations. One is centered at *δ*_1_ ≈ 85° and *δ*_2_ ≈ 150°, consistent with a C2^′^-endo conformational state. The second is located near *δ*_1_ ≈ 85° and *δ*_2_ ≈ 85°, where both nucleotides adopt a C3^′^-endo pucker. Notably, this model successfully samples both minima, demonstrating its improved description of the underlying conformational landscape. Although the *δ*_1_ ≈ 85° and *δ*_2_ ≈ 85° dihedral population is less pronounced, the overall agreement indicates that RNA-DFT captures the essential features of the reference distribution, even in the absence of explicit solvent or long-range stacking interactions.

To evaluate the structural accuracy of the different ML force fields, we further analyzed the probability density distributions of the four additional RNA backbone torsion angles (*γ, ε, ζ*, and *β*) for the ApA dimer. As shown in Fig. S9, these angles define the local geometry of the phosphodiester backbone and are subject to strict conformational constraints that characterize canonical A-form ^73,74^. In A-form conformation, the *γ* angle is essential for maintaining the C3’-endo sugar pucker and the overarching A-form geometry, which requires a gauche-plus conformation centered near +60° ^73,74^. Both TREMD and our model exhibited sharp, well-defined peaks at this value, a trend also preserved by SO3LR and RNA-DFT. The *β* torsion typically populates the trans region around −150°. This feature is well captured by both TREMD and our model, whereas SO3LR exhibits a broader distribution, consistent with increased backbone flexibility reported along the *δ* angles. Canonical values for the *ε* and *?* torsion angles around −65° ^74^ are accurately reproduced by all ML models.

We computed the Hellinger distance (see Eq. 2) between the one-dimensional distributions of each backbone and glycosidic torsion angles, comparing them with the TREMD reference distributions (see Fig. 6). This metric allows for a direct comparison of how closely each model approximates the conformational ensemble observed in the reference data. Lower Hellinger distances indicate higher similarity. The RNA-DFT consistently achieves the lowest Hellinger distances for the *χ*, and *δ, γ* and *ε* torsion angles, which are among the most structurally relevant angles for RNA conformational description. In particular, the *δ* angle governs the sugar pucker conformation (C3^′^-endo versus C2^′^-endo), while the *χ* angle controls the base orientation (anti-syn conformations). Accurate reproduction of these angles is critical for modeling base stacking and helicity in RNA dimers. The *ε* angle, associated with backbone flexibility and phosphate positioning, is also captured with high accuracy by the RNA-DFT model relative to the reference TREMD data, highlighting its suitability for sampling RNA conformations. The RNA-TB model performs comparably better than the general-purpose SO3LR model for several torsion angles, especially *δ* and *χ*, but shows higher deviations for other backbone angles.

**Figure 6.**
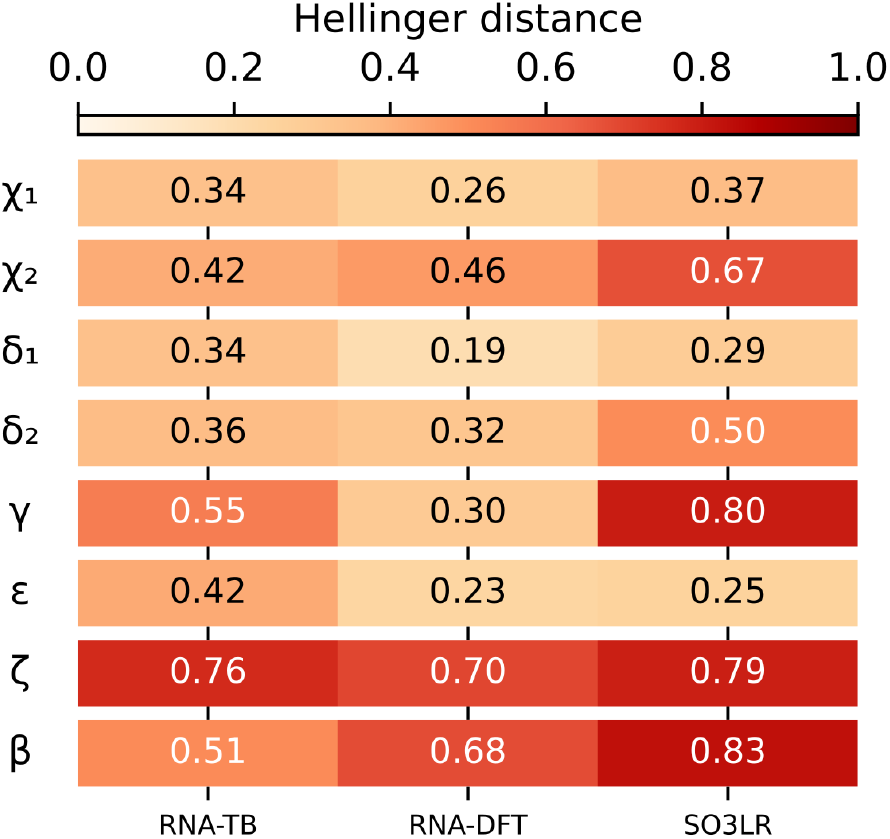
Hellinger distances between the probability distributions of each torsional angle computed from different models and the reference TREMD simulation. Lower values indicate higher similarity to the TREMD reference.

## 4 Conclusions

In conclusion, we assessed the performance of quantum-accurate ML potentials in exploring conformational transitions in RNA dimers. As a representative case, we selected the adenine–adenine dinucleoside monophosphate (ApA) system and constructed a comprehensive QM dataset based on exhaustive conformational sampling obtained from TREMD simulations. This approach yielded a diverse ensemble of stacked and unstacked conformations, which we classified into six clusters: A-form, inverted, ladder, anti-ladder, sheared, and unstacked. To evaluate how the electronic description of interatomic interactions in solvated ApA influences ML performance, we generated QM datasets using two electronic-structure methods: the semi-empirical third-order density-functional tight-binding method (DFTB3) and density functional theory (DFT) with the PBE0 hybrid functional. Both methods were complemented by a many-body dispersion (MBD) treatment to account for van der Waals interactions, which are essential for accurately describing long-range electronic effects during structural transitions in condensed-phase systems.

Our analysis of the two QM datasets revealed fundamental differences in how these electronic-structure methods describe the ApA conformational landscape. In particular, PBE0+MBD exhibits a broader atomization-energy range and larger MBD contributions than DFTB3+MBD, indicating improved energetic discrimination among distinct conformations. Further analysis of atomic charge fluctuations along the reference TREMD trajectory highlights the importance of capturing charge redistribution when developing RNA force fields—an effect typically neglected in classical force-field parameterizations but inherently included in ML potentials trained on QM energies and forces. Consistent with these observations, the ML potential trained on DFT data (RNA-DFT model) reproduces stacking fractions and A-form populations more accurately relative to the TREMD reference than the RNA-TB model. General-purpose ML models, such as MACE-OFF24 and SO3LR, also predict high stacking fractions; however, their A-form populations deviate substantially from the TREMD results. Moreover, analysis of conformational transitions shows that, unlike the RNA-DFT model, the SO3LR model preferentially samples direct transitions between stacked clusters (*e*.*g*., A-form, anti-ladder, and inverted), suggesting an incomplete description of the underlying transition pathways. A detailed analysis of the A-form in terms of the *χ* and *φ* dihedral angles further clarifies these differences. The SO3LR model deviates primarily in the *φ* distributions, overestimating conformational flexibility relative to the TREMD reference, whereas the RNA-DFT model exhibits substantially better agreement. This trend is quantitatively confirmed by statistical analysis using the Hellinger distance, with the overall agreement ranked as follows: RNA-DFT > RNA-TB > SO3LR.

Overall, these findings highlight two central challenges in exploring RNA conformational landscapes with quantum-accurate ML potentials: (i) the scarcity of QM datasets that exhaustively sample the conformations of RNA building blocks and (ii) the limitations of current state-of-the-art ML potentials in capturing complex structural transitions in biological systems, even for a relatively small system such as the ApA dimer. Addressing these challenges will enable the development of more accurate and transferable ML potentials capable of reliably describing structural transitions and predicting RNA conformations and relative energetics. This need is also motivated by the rapidly increasing number of non-coding RNA sequences requiring a 3D structural characterization. Furthermore, the generated datasets may be interpolated by the aforementioned ML potentials and integrated with fast MD sampling. In this manner, a combined framework could be used to reweight atomistic molecular simulations to locally correct force-field inaccuracies, provided that the structure is decomposed into smaller fragments. In a similar way, the accuracy of the ML potentials could be assessed with respect to NMR data^75,76^ or thermodynamic quantities^77,78^. We therefore expect that the insights presented here will contribute to the design of ML-assisted computational frameworks for the next generation of quantum-accurate RNA force fields.

## Data availability

All datasets, running examples, and trained ML models presented in this work are available on Zenodo under the DOI 10.5281/zenodo.18714738.

## Acknowledgements

LMS and GC gratefully acknowledge the funding by the German Research Foundation (DFG) under the Cluster of Excellence CeTI: Centre for Tactile Internet with Human-in-the-Loop (EXC 2050/2, Project ID 390696704), Cluster of Excellence REC^2^: Responsible Electronics in the Climate Change Era (EXC 3035), and Cluster of Excellence CARE: Climate-Neutral And Resource-Efficient Construction (EXC 3115, Project ID 533767731). SP was supported by the Fondecyt Regular project No. 1231071 from ANID, Chile and Centro Ciencia & Vida, FB210008, Financiamiento Basal para Centros Científicos y Tecnológicos de Excelencia de ANID. ABP acknowledges financial support from the National Science Center, Poland, under grant 2022/45/B/NZ1/02519. LMS and MTN are grateful to the Research Experience for Peruvian Undergraduates (REPU) program for its organizational support. We thank the Center for Information Services and High-Performance Computing (ZIH) at TU Dresden and the Polish high-performance computing infrastructure PLGrid (HPC Center: ACK Cyfronet AGH, computational grant no. PLG/2024/017332 and PLG/2025/018510) for providing the computational resources and technical support.

## Author contributions

The work was initially conceived by LMS, SP, and ABP, and designed with contributions from MTN and LFCV. LMS and MTN developed the QM datasets and the ML potentials and analyzed the model performance. SP performed the PMD and TREMD simulations. GEOR participated in scientific discussions. LFCV and ABP performed clustering analysis and validated the ML potentials in MD simulations and further structural analysis. LMS, GC, SP, and ABP supervised and revised all stages of the work. All authors discussed the results and contributed to the final manuscript.

