## Supplementary Information for "Exploring Conformational Transitions of RNA Dimers via Machine Learning Potentials"

### 1 RNA simulation details

Here we will enumerate and describe the data generated for the ML analysis.

#### 1.1 ApA-TREMD

It corresponds to the Temperature-Replica Exchange MD simulation of a Adenine-Adenine dinucleotide monophosphate. That is, two nucleotides connected through the backbone where the first one does not have a phosphate group. In consequence, a single positive ion must be included to keep the neutrality of the whole system. The DMNP is solvated in 1080 water molecules and a Sodium Ion in a cubic box of length $L = 32.1719$Å using periodic boundary conditions. The initial condition was taken from an A-form duplex, and then energy-minimized with a steepest-descent algorithm, to be relaxed with a short NpT simulation for 1 ns.

Following the notation of the work of Hayatshahi et al.[1], the force field employed was ff14 (Amber99 with parmbsc9 and $\chi$OL corrections) with a TIP3P water model. Following the same reference, we used a Temperature-Replica Exchange Molecular Dynamics simulation for 500 ns, which was reported to sample the conformation space satisfactorily. The MD was integrated using the leap-frog method with a timestep of 0.002 ns, while temperature was kept constant with a velocity rescaling with a stochastic term thermostat with a constant of 0.1 ps. The temperatures were 280, 285.8, 291.7, 297.7, 303.9, 310.1, 316.6, 323.1, 329.8, 336.6, 343.5, 350.6, 357.9, 365.3, 372.8, 380.5, 388.4 and 396.4 K where the exchange acceptance was between 20 and 30%. We checked that the temperature space was thoroughly and frequently visited by each replica. Finally, we selected the simulation at T=297.715K .

##### *1.1.1 Data files*

For the selected simulation, we have deployed the trajectory in pdb and xyz formats, composed of snapshots saved every 50 ps, giving a total of 10000 frames. The coordinates of the atoms were centered around the group of atoms composing the RNA molecule on each timestep. The rest of the files are in xvg format, which is an ascii file where the data is stored in lines for each timestep. The data is extended along columns, whose content is detailed in the first commented lines in the file, starting with the characters "`@`" or "`#`".

An offline analysis of the forces was performed on the selected simulation. The file `cropped_force.xvg` contains the force on each atom, which are numbered according the pdb file. Each line contains the timestep in each column, followed by the three coordinates of the total force on every different atom. The first lines of the datafile, which begin with the character "`@`", detail this correspondence.

The energies of the whole system are also calculated, keeping track of different contributions. Taking into account the energy terms of the AMBER force field, it makes sense to decompose the energy between the groups of RNA and non-RNA atoms. The second is composed by the water molecules and the Na ion. These groups are labeled as `RNA` and `IWAT`, respectively. The energies are reported for the interactions between `RNA-RNA`, `RNA-IWAT` and IWAT-IWAT. Given the treatment of the electrostatic interactions, and additional term which is not possible to decompose under this approach is the Coulomb-reciprocal, which is also included for completeness. In order to provide more detail in the contributions of each of these terms, the xvg files are divided into bonded (only for intra-RNA interactions) and non-bonded, which have terms of Lennard-Jones and Coulomb in different columns on each timestep. The lines in the header of the file specify which interaction corresponds to each column. Nevertheless, the sum over the columns of an energy file, excluding the time, yield the total energy for bonded or non-bonded cases.

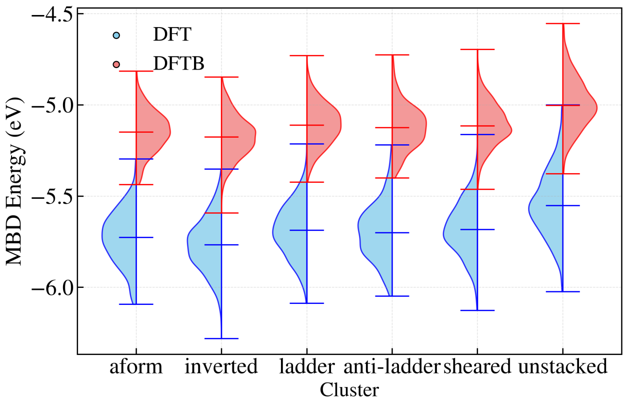


**Figure S1.** Distributions of the MBD energies obtained with DFT and DFTB for ApA structures containing 194 atoms. We have divided them per conformational cluster.

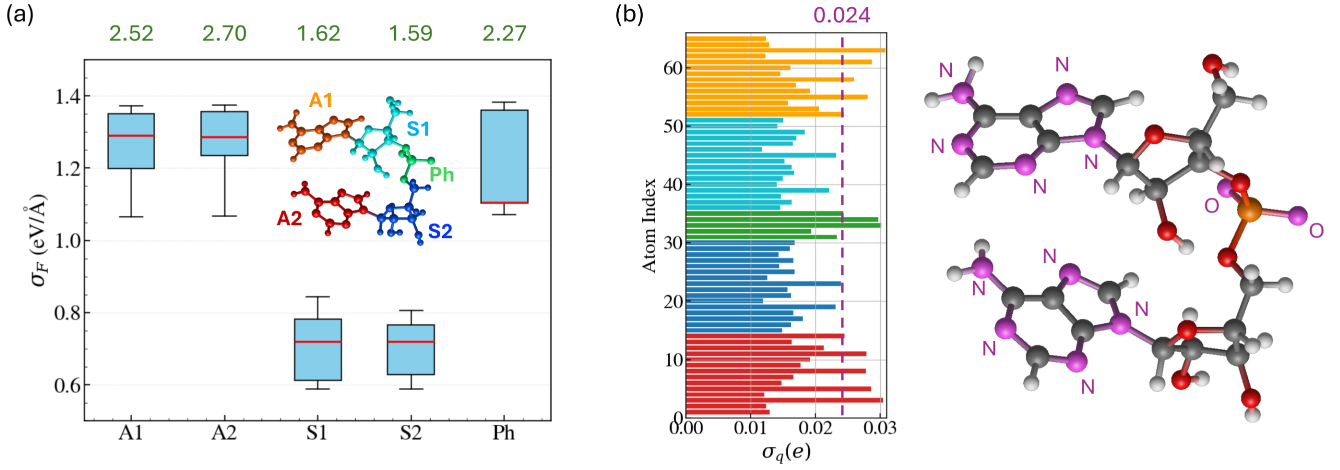


**Figure S2.** (a) Box plots of the standard deviation of atomic forces, $\sigma_F$, computed with DFTB for the ApA building blocks: adenine 1 (A1), adenine 2 (A2), sugar 1 (S1), sugar 2 (S2), and phosphate (Ph). Mean force values for each group are also shown on the top of box plots. (b) We show the variance of atomic charges, $\sigma_q$, computed with DFTB for all atoms in ApA dimer. The atom index has been ordered with respect to the molecular groups defined in panel (a). In the inserted structure, we have highlighted in violet the atoms with $\sigma_q > 0.024e$.

###### *1.1.2 Stability of ApA centroids in MD simulation using MLPs*

A visual and quantitative evaluation of the degree to which the sampled conformers from simulations resemble the five reference stacking conformations is provided by the structural overlays in Figure S4. The RMSD values reported correspond to the frame with the lowest deviation from each reference structure, providing insight into the accuracy of sampled conformations. In general, the inverted conformation showed the best structural agreement across these four models, with RMSD values consistently around 0.12-0.24 Å. This means that this conformation was not only frequently sampled, as indicated by the population data, likely due to its stable stacking conformation. The A-form conformation resulted in lower RMSD values (approx. 0.15 Å) in SO3LR follower by both models RNA-TB and RNA-DFT with 0.52 Å) and m4-8 (0.25 Å) respectively. The ladder and sheared conformations showed greater structural variability across the models and sampling schemes. The RMSD values for ladder was 1.30 Å for RNA-TB, 1.33 Å for RNA-DFT and 1.41 Å for SO3LR . The sheared conformation showed a comparable pattern, with RMSD values exceeding 1 Å for both RNA models. The anti-ladder conformation showed moderate agreement with the reference in all conditions. RMSD values were consistently in the range of 0.23-0.45 Å, indicating that this conformation is sampled with consistency and may represent a recurrent intermediate state between canonical and unstacked forms. The SO3LR sampled significantly very close conformations of the ApA system. Either means that SO3LR over-fitted the A-form conformation or lacked sensitivity to capture alternative stacking arrangements.

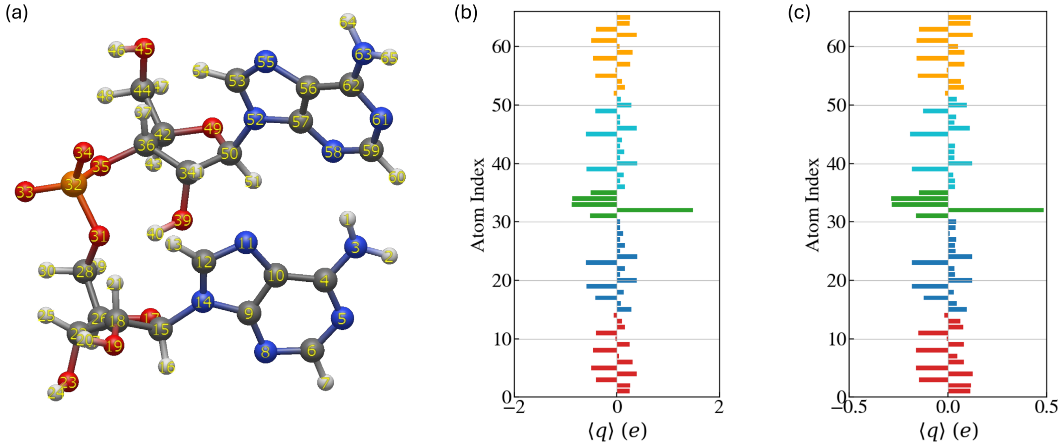


**Figure S3.** (a) ApA structure displaying the indices used for the analysis of atomic charges, $q$, in the Fig. 2 of the main text and Fig. S2 of the SI. We show the mean value of atomic charges, $\langle q \rangle$, computed by (b) DFT and (c) DFTB.

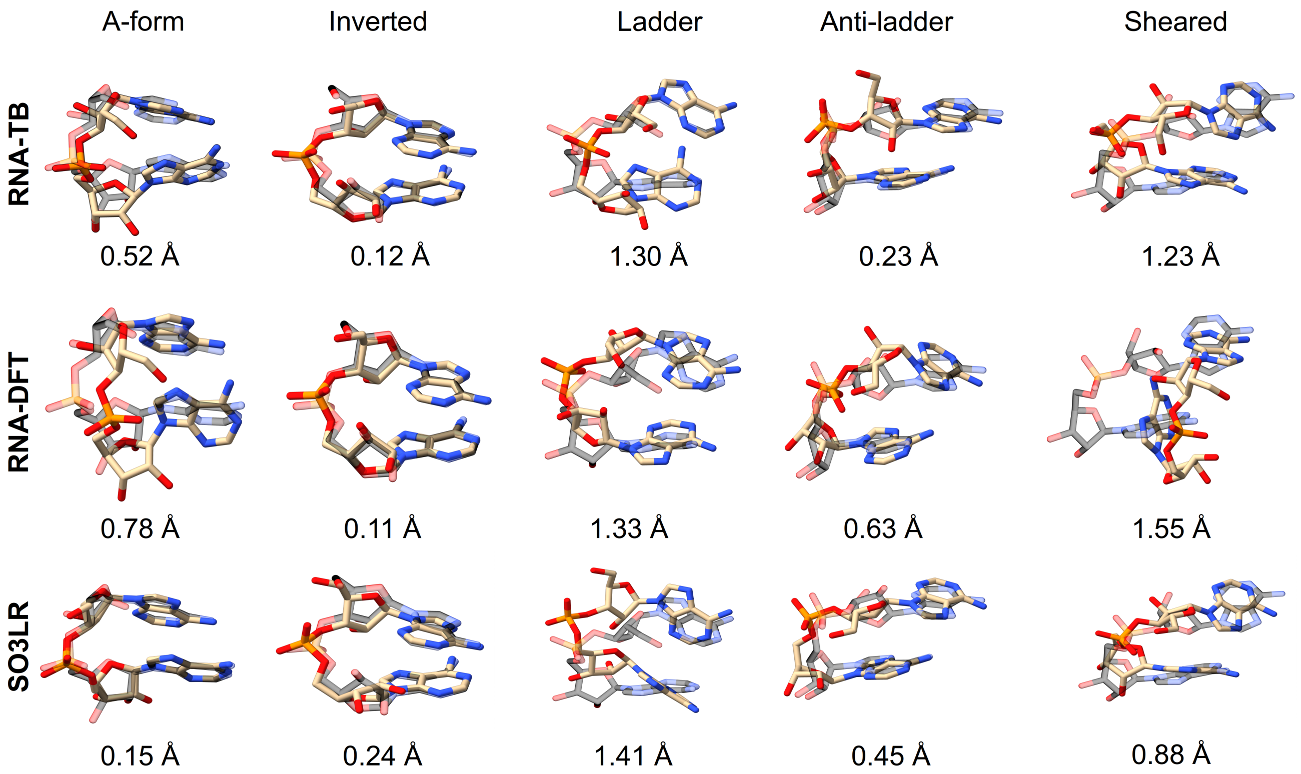


**Figure S4.** Overlay of representative ApA RNA dimer conformations with their corresponding reference structures for five stacked geometries: A-form, inverted, ladder, anti-ladder, and sheared. Transparent models indicate the reference structures, while opaque models represent the simulation frame with the lowest RMSD to each reference. Rows A and C correspond to the lowest RMSD conformers from PMD simulations of models m4-7 and m4-8, respectively. Rows B and D show the corresponding conformers from TREMD simulations for the same models. Row E presents the closest conformers obtained from SO3LR simulations, and Row F shows those from MACE-OFF-24 predictions. Reported RMSD values (in Å) between representative frames and their respective reference conformations.

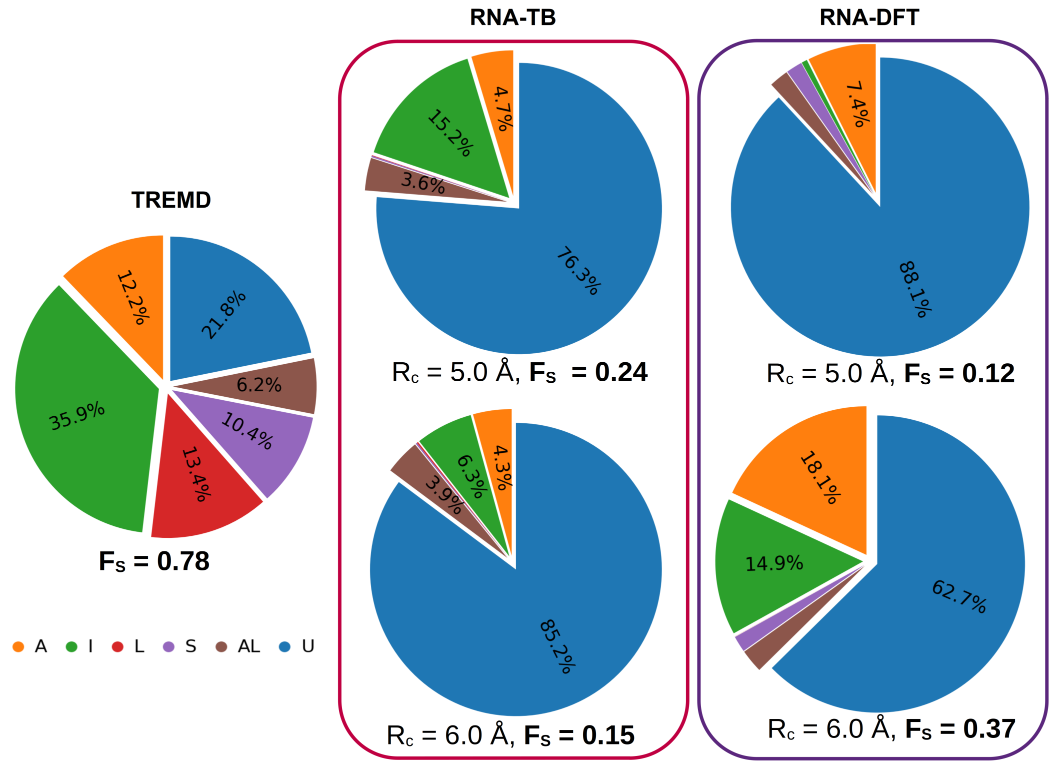


**Figure S5.** Comparison of stacking conformational ensembles and transition behavior for ApA dimers across models. Pie charts depict the relative populations of six conformational classes: A-form (A), inverted (I), ladder (L), anti-ladder (AL), sheared (S), and unstacked (U).

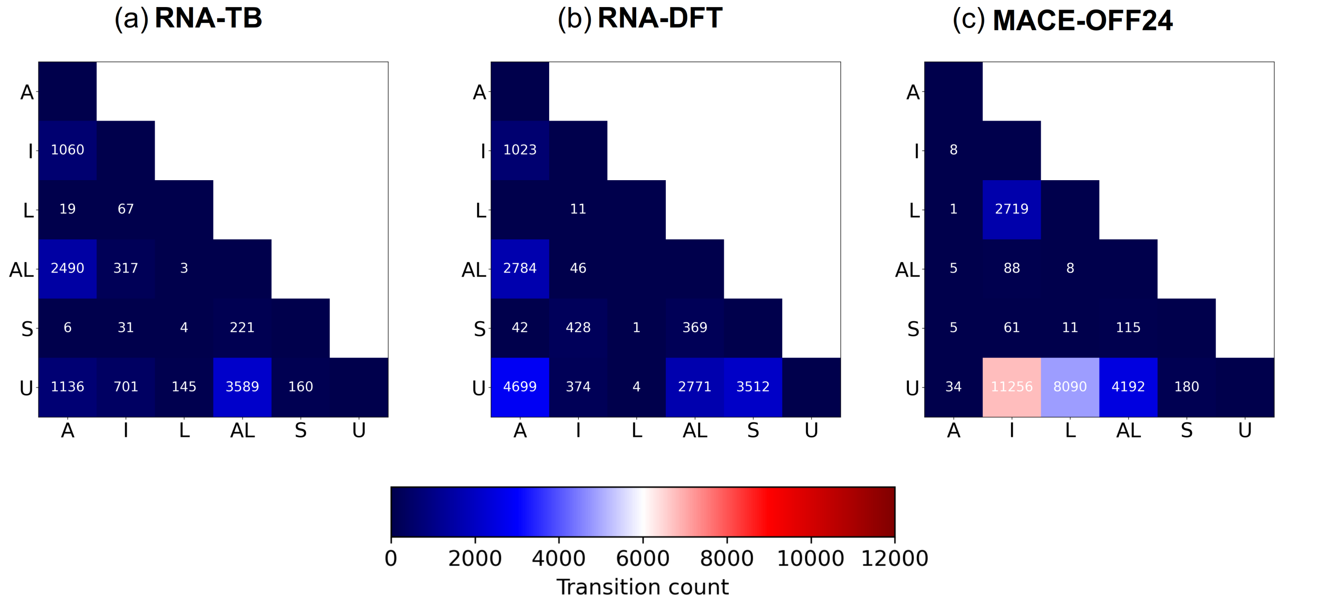


**Figure S6.** Transition behavior for ApA dimers across models. Stacking transition matrices show the number of transitions between stacking states for each model. Each panel corresponds to a different method, with transitions accumulated from three 10-ns simulations per model (sampled every 10 fs, yielding $10^6$ frames). Cutoff radius of $R_c = 6$ Å for RNA-TB and $R_c = 5$ Å for RNA-DFT.

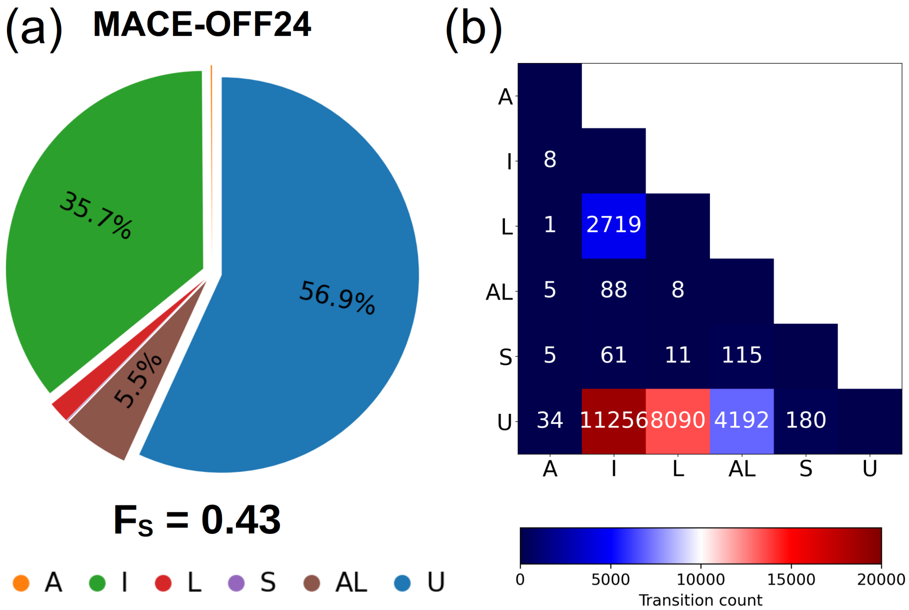


**Figure S7.** Comparison of stacking conformational ensembles and transition behavior for ApA dimers in MACE-OFF24 model. (A) Pie chart depicts the relative populations of six conformational classes: A-form (A), inverted (I), ladder (L), anti-ladder (AL), sheared (S), and unstacked (U). (B) Stacking transition matrix shows the number of transitions between stacking states).

**Table S1.** Atomic nomenclature of RNA bases and sugar-phosphate backbone

| Component | Atomic nomenclature |
|---|---|
| Base | N1, C2, N3, C4, C5, C6, N7, C8, N9 |
| Backbone | P, OP1, OP2, O5', C5', C4', O4', C3', O3', C2', O2', C1' |

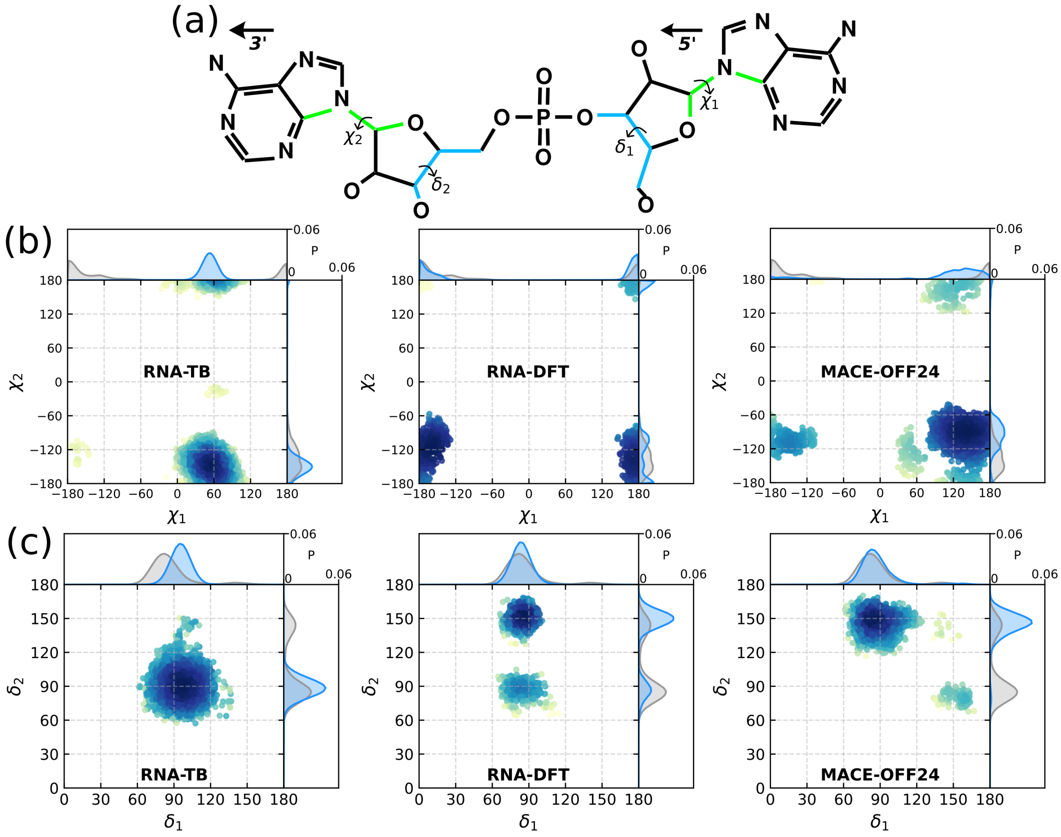


**Figure S8.** (A) Chemical structure of the ApA RNA dimer showing the definition of the $\chi$ and $\delta$ torsion angles for both nucleotides. (B) Two-dimensional distributions of $\chi_1$ and $\chi_2$ angles sampled during simulations using the RNA-TB ($R_c = 6$ Å), RNA-DFT ($R_c = 5$ Å), and MACE-OFF24 models. (C) Two-dimensional distributions of $\delta_1$ and $\delta_2$ angles for the same models. The color scale represents the normalized probability of sampling each pair of torsional angles. The reference TREMD distributions are shown in gray color.

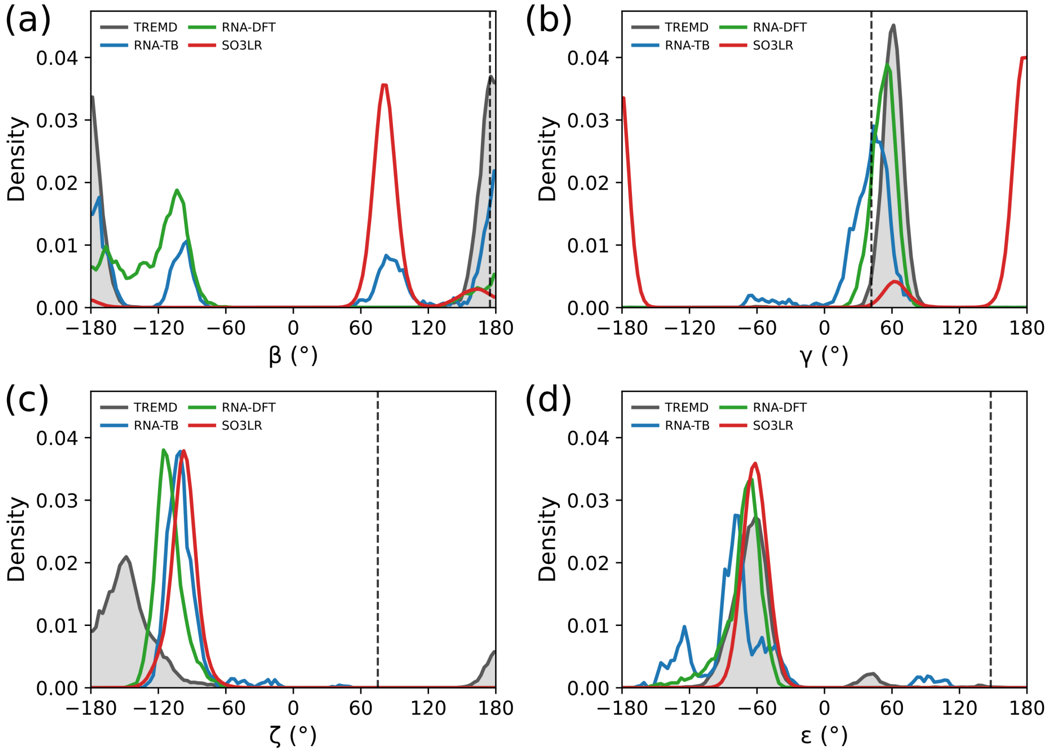


**Figure S9.** (A) Chemical structure of the ApA RNA dimer showing the definition of the $\beta$, $\gamma$, $\zeta$ and $\varepsilon$ torsion angles. (B) Single distributions for the reference TREMD data and the RNA-TB ($R_c = 5$ Å), RNA-DFT ($R_c = 6$ Å), SOL3R and MACE-OFF24 models. Vertical dashe lines indicate the experimental values for A-form[2].
